# SNAP-tag and HaloTag fused proteins for HaSX8-inducible control over synthetic biological functions in engineered mammalian cells

**DOI:** 10.1101/2022.08.12.503781

**Authors:** Hannah L. Dotson, John T. Ngo

## Abstract

Drug-inducible systems allow biological processes to be regulated through the administration of exogenous chemical inducers. Such methods can be used to study native biological activities, or to control synthetically engineered ones, with temporal and dose-dependent control. However, the number of existing drug-inducible systems is limited, and there remains a need for synthetic biology components that can be combined with the existing toolset and regulated with independent and orthogonal control. Here, we describe new cell engineering components that can be regulated via a heterodimerization of SNAP-tag and HaloTag domains using a selective small molecule crosslinker termed “HaXS8.” The construction and validation of multiple HaXS8-sensitive components are described, including systems for regulating transcription, Cre recombinase activity, and caspase-9 activity in mammalian cells. The systems elaborate the ability to control gene expression, DNA recombination, and apoptosis in cell engineered systems.

## INTRODUCTION

One of the essential methods that nature uses to control cellular behavior is regulation of protein interactions. As with many natural processes, synthetic biology seeks to replicate and expand on this behavior, both to understand existing natural systems and to drive specific behavior in synthetic systems. One technique to induce protein localization involves the usage of “bifunctional” small molecules, which can interact with two proteins simultaneously, in order to induce proximity between fused domains^1^. These chemically-inducible dimerization (CID) systems have been widely used to investigate native biological systems such as signaling pathways^2^, and have also been used to regulate a wide variety of designed biological behavior, including cell signaling^3^, protein degradation^4,5^, protein splicing^3,6^, apoptosis^7^, construction of logic gates^8^, gene expression^9,10^, and gene editing^11^.

Perhaps the most commonly used CID platform is the rapamycin-induced FKBP/FRB system ^12^. In this system, the binding of rapamycin to FKBP12 produces a protein-ligand complex that is then recognized by FRB. In cells, proximity between sequences that are fused to FKBP and FRB can be induced via rapamycin and its analogs, and this approach has been used to precisely control cell signaling^3^, protein degradation^4,5^, protein splicing^6^, and apoptosis^7^, as well as many other biological processes.

A variety of other CID methods adapting natural systems have been more recently developed, including the abscisic acid(ABA)-inducible ABI/PYL^13^, Mandipropamid-inducible ABI/PYR^Mandi14^, and gibberellin(GIB)-inducible GAI/GID1^8^ systems. Several of these systems have been demonstrated to be orthogonal to existing CID systems, allowing for the design of logic gates involving multiple CIDs^8,15,16^. Engineered CID systems have also been constructed, such as the antibody-based chemically induced dimerizer (abCID)^17^ system and the nanobody-based combinatorial binders-enabled selection of CID (COMBINES-CID)^18^ system, both of which demonstrate general methods to create new CID systems. As these systems continue to be developed, versions with additional features, such as spatiotemporal control via light-activated methods, have been created^19^.

Despite these advances, development of novel CID systems and new applications of existing CID systems remain an area of interest. Additional orthogonal CID systems can allow for more complex, multi-input behavior^8^, and expanding on existing CID systems provides new methods of controlling different protein behavior. Inspired by the success of these existing CID systems, we sought to extend the CID system “toolbox”. Given current work in the development of bifunctional, proximity-inducing small molecules^1^, we took specific interest in a chemical dimerizer of HaloTag and SNAP-tag, HaXS8.

SNAP-tag and HaloTag are self-labeling protein tags that are generally used with corresponding probes for imaging, among a variety of other uses. SNAP-tag, derived from human O6-alkylguanine DNA alkyltransferase ^20–22^, reacts with benzylguanine derivatives and was developed as a method to covalently label fusion proteins *in vivo*. It has since been used in conjunction with functional probes for a wide variety of purposes, including fluorescence imaging^23^, immobilization^24^ and localization^25^ of proteins, and optochemical control of systems^26^. HaloTag^27^, derived from a bacterial haloalkane dehalogenase, reacts with a synthetic ligand containing a chloroalkane linker. Usages of HaloTag include control of protein degradation via the HaloPROTAC system^28^, protein purification from bacterial^29^ and mammalian^30^ cells, and fluorescence imaging^27,31^. In previous work, the SNAP-tag and HaloTag domains have been used together to create a heterodimerization system, via a small bifunctional molecule ligand HaXS8^32^ (**Figure 1a, b**). HaXS8 consists of an O6-benzylguanine which acts as a substrate for SNAP-tag, a chloroalkane which acts as a substrate for HaloTag, and a core module which renders the molecule cell permeable^32^. The HaXS8 heterodimerization system was shown to have the ability to force protein-protein interactions, and was specifically used to induce protein translocation to different cellular compartments and activate the PI3K/mTOR pathway, which generally cannot be activated with rapalog CID systems without crosstalk due to rapamycin’s interaction with mTOR^32^.

**Figure 1.**
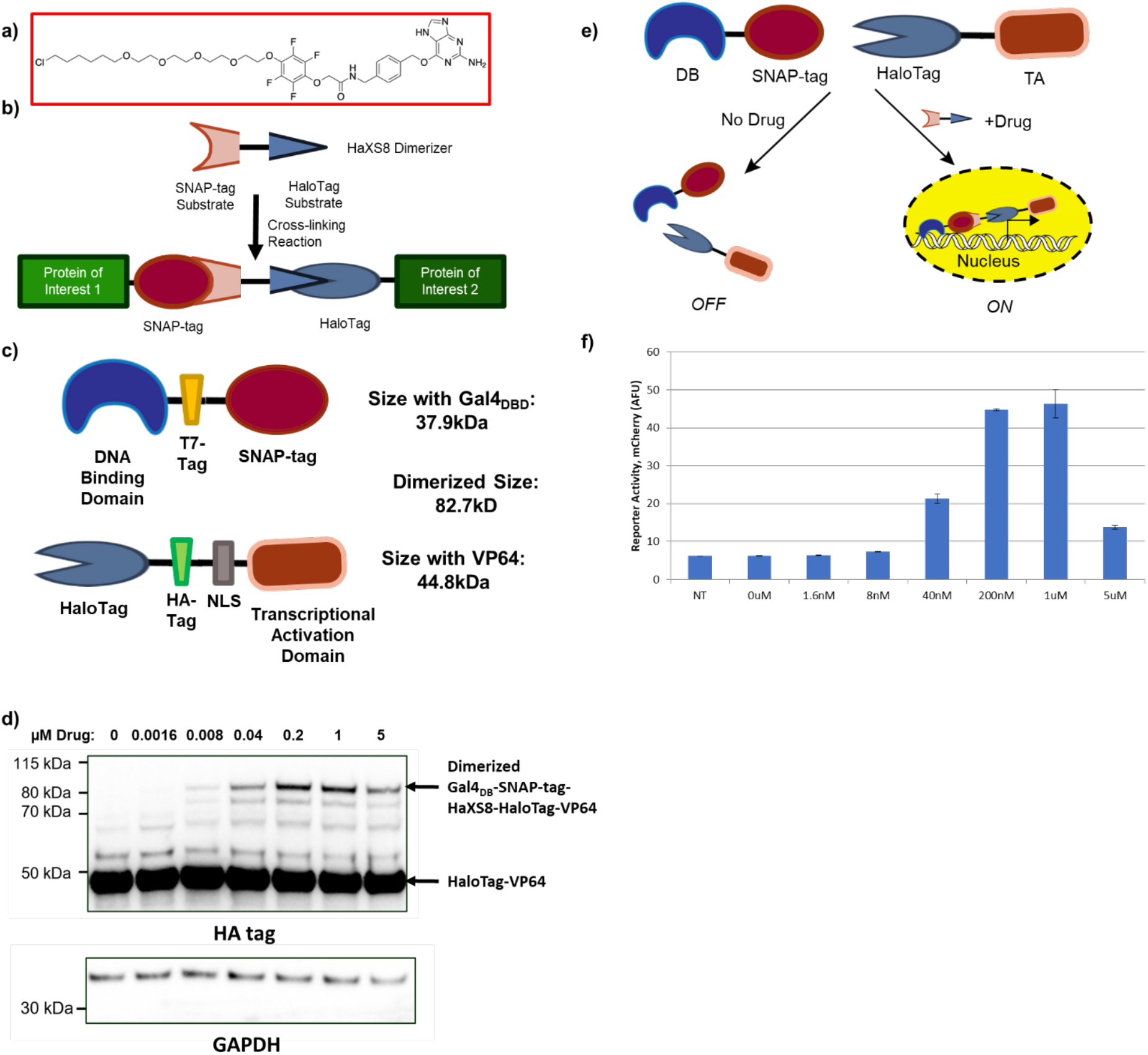
Design of a HaXS8-inducible TF system. **a)** Chemical structure of the HaXS8 molecule. **b)** HaXS8 consists of a SNAP-tag substrate and a HaloTag substrate connected by a linking module. HaXS8 is able to heterodimerize proteins of interest fused to SNAP-tags and HaloTags. **c)** Schematic showing the structure of the HaXS8-inducible TF system, as well as tag locations for immunoblotting. **d)** Western blot showing accumulation of full-length, dimerized Gal4_DB_-SNAP-tag-HaXS8-HaloTag-VP64 (anti-HA; 82.7 kDa) in response to HaXS8. Undimerized HaloTag-VP64 (anti-HA; 44.8 kDa) is also observed. GAPDH (anti-GAPDH; 37 kDa) was used as a loading control. **e)** When no drug is present, DB-SNAP (DNA-Binding Domain-SNAP-tag) and Halo-TA (HaloTag-Transcriptional Activation Domain) remain separate, and do not act as a functional TF. When HaXS8 is added, the two halves are able to heterodimerize, resulting in a functional TF. This functional TF allows transcription of a target gene that corresponds with concentration of drug. **f)** H2B-mCherry fluorescence as determined by flow cytometry in Hek293FT reporter cells (UAS H2B-mCherry) transiently transfected with plasmids encoding Gal4_DB_-SNAP-tag and HaloTag-VP64 at a 1:1 ratio and treated with varying concentrations of HaXS8. Geometric means are displayed as mean ± s.d., as determined by three transfected cell cultures.

In addition to the successful applications already noted, the HaXS8 system had a variety of other useful properties. In terms of implementation, HaXS8 was commercially available, making the small molecule itself simple to obtain. Similarly, SNAP-tag and HaloTag are commonly used domains, meaning that obtaining their sequences from existing plasmids present in the lab or through suppliers (including the non-profit repository AddGene) was also straightforward. Furthermore, the widespread utility of SNAP-tag and HaloTag also mean that new variations and ligands are continually being engineered. Recent examples of such work include Photo-SNAP-tag^33^, a light-regulated split SNAP-tag, and HaloTag variants that lead to different brightness and fluorescence lifetimes following single fluorophore ligation^31^. Exchangeable HaloTag ligands^34^ and assembly-regulated SNAP-tag probes^35^ have also recently been described. In this work, we describe additional SNAP-tag and HaloTag utilities, which are used in combination together with HaXS8 to achieve precise control over synthetic biology activities including transcription, recombination, and triggered-cell death.

## RESULTS

We first wanted to demonstrate that the SNAP-tag/HaloTag heterodimerizer system with HaXS8 could be used to control activity of a transcription factor (TF) in a drug-inducible manner. We constructed a split-TF system consisting of two constitutively-expressing plasmids, one expressing a DNA-binding Domain (DB) and SNAP-tag fusion and the other expressing a HaloTag and transcriptional activator (TA) fusion (**Figure 1c**). Immunoblotting of transfected mammalian cells showed drug-dependent formation of the cross-linked Gal4_DB_-SNAP-tag /HaloTag-VP64 species (**Figure 1d, Supplemental Figure 1**; 82.7 kDa), with detectable levels of the covalent heterodimer forming within 30 minutes of drug exposure (**Supplemental Figure 2**). The presence of the cross-linked heterodimer was not observed in drug untreated control cells, and we note that formation of the dimeric product was diminished at high drug concentrations (5 μM; **Figure 1d**), likely due to the oversaturation of the individual Gal4_DB_-SNAP-tag and HaloTag-VP64 halves with HaXS8.

Having confirmed the ability to induce dimerization of our protein components, we next tested the activity of the cross-linked TF product via a cell-based transcriptional reporter assay (**Figure 1e**). Here, we expressed Gal4_DB_-SNAP-tag and HaloTag-VP64 at a 1:1 ratio in HEK293FT cells containing a genetic reporter construct (UAS H2B-mCherry) and quantified the expression of the reporter following overnight incubation in the presence or absence of HasX8. Quantification of reporter expression levels by flow cytometry confirmed the transcriptional activity of the expressed split-TF (**Figure 1f**), which facilitated reporter expression in a HaXS8 dose dependent manner. Notably, reporter expression levels in response to HasX8 mirrored the dose-dependence of heterodimer formation observed in our immunoblot analyses, with a maximal reporter response observed in the range of 200 nM to 1 μM and with diminished H2B-mCherry expression yields at elevated concentrations. Furthermore, reporter expression in a drug untreated control was comparable to a nontransfected control (**Figure 1f**). Consistent with these analyses, expression of Gal4_DB_-SNAP-tag and HaloTag-VP64 individually (without their corresponding dimerization partner) did not result in reporter activation in either HasX8- or drug-untreated conditions **(Supplemental Figure 3**). Together, these results confirm that our HaXS8-inducible split-TF system is able to drive transcription in the presence of drug, but not in the absence of drug or when only one half of the system is expressed.

Having confirmed the ability to achieve Gal4-mediated transcriptional control, we next tested the versatility of the HaSX8 strategy by installing the same mode of regulation over TF components targeting distinct and orthogonal promoters. Here, we exploited the modularity of our design by substituting our initial DB domain (Gal4_DB_) with QF_DB_, which is derived from the Q regulatory system of *Neurospora qa* and binds the QUAS promoter^36,37^. Fusion of QF_DB_ to the SNAP domain generated QFDB-SNAP-tag (**Figure 2a**), which we tested in combination with HaloTag-VP64 using a QUAS H2B-citrine reporter construct. Satisfyingly, analyses by flow cytometry confirmed portability of HaXS8-inducible transcriptional regulation, with cells exhibiting drug-dependent reporter expression responses similar to those that were observed via our Gal4-based approach above (**Figure 2b**).

**Figure 2.**
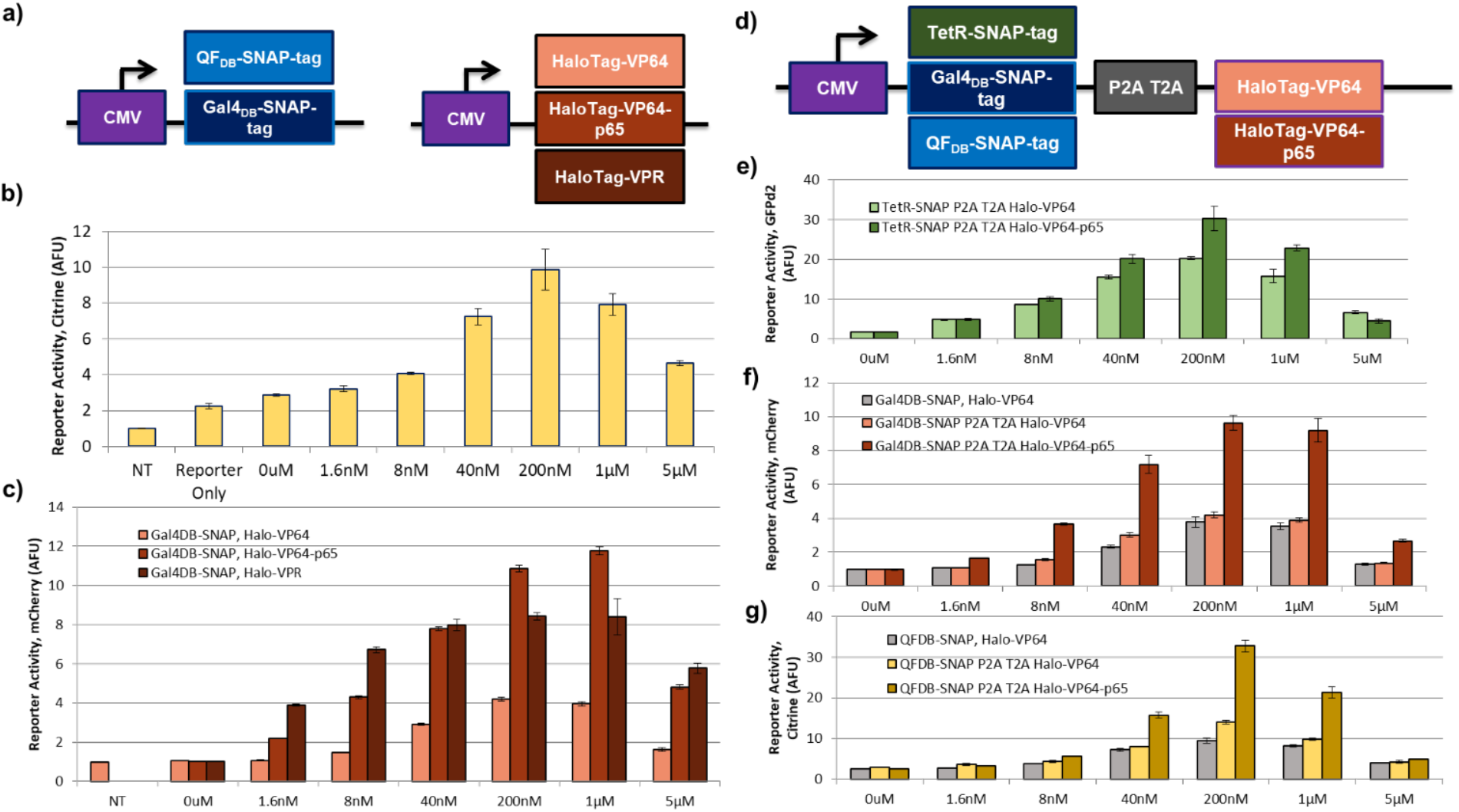
Designs with varying DB and TA domains for a HaXS8-inducible “Turn-on” system. **a)** Schematic showing the plasmid schematics for the two halves of the HaXS8-inducible TF: DB-SNAP (DNA-Binding Domain-SNAP-tag) and (Halo-TA) HaloTag-Transcriptional Activation Domain. **b)** H2B-citrine fluorescence as determined by flow cytometry in Hek293FT cells transiently transfected with a QUAS H2B-citrine reporter and plasmids encoding QFDB-SNAP-tag and HaloTag-VP64 at a 1:1 ratio, treated with varying concentrations of HaXS8. **c)** H2B-mCherry fluorescence as determined by flow cytometry in Hek293FT reporter cells (UAS H2B-mCherry) transiently transfected with plasmids encoding Gal4_DB_-SNAP-tag and varying HaloTag-TAs at a 1:1 ratio and treated with varying concentrations of HaXS8. **d) *Single vector design for a HaXS8-inducible “Turn-on” system*.** Schematic showing the design of a single vector encoding both halves of the HaXS8-inducible “turn-on” system. Both halves of the HaXS8-inducible TF, DB-SNAP (DNA-Binding Domain-SNAP-tag) and Halo-TA (HaloTag-Transcriptional Activation Domain), are encoded on one plasmid, separated by a P2A T2A sequence. **e)** GFPd2 fluorescence as determined by flow cytometry in Hek293FT reporter cells (TRE3G GFPd2) transiently transfected with a plasmid encoding either TetR-SNAP-tag P2A T2A HaloTag-VP64 or TetRSNAP-tag P2A T2A HaloTag-VP64-p65, treated with varying concentrations of HaXS8. **f)** H2B-mCherry fluorescence as determined by flow cytometry in Hek293FT reporter cells (UAS H2B-mCherry) transiently transfected with a plasmid encoding either Gal4_DB_-SNAP-tag P2A T2A HaloTag-VP64 or Gal4_DB_-SNAP-tag P2A T2A HaloTag-VP64-p65, treated with varying concentrations of HaXS8. For comparison, Hek293FT reporter cells (UAS H2B-mCherry) transiently transfected with plasmids encoding Gal4_DB_-SNAP-tag and HaloTag-VP64 at a 1:1 ratio, treated with varying concentrations of HaXS8, are also shown. **g)** H2B-citrine fluorescence as determined by flow cytometry in Hek293FT cells transiently transfected with a QUAS H2B-citrine reporter and a plasmid encoding either QFDB-SNAP-tag P2A T2A HaloTag-VP64 or QFDB-SNAP-tag P2A T2A HaloTag-VP64-p65, treated with varying concentrations of HaXS8. For comparison, Hek293FT cells transiently transfected with a QUAS H2B-citrine reporter and plasmids encoding QFDB-SNAP-tag and HaloTag-VP64 at a 1:1 ratio, treated with varying concentrations of HaXS8, are also shown. Geometric means are displayed as mean ± s.d., as determined by three transfected cell cultures. Data is normalized such that NT samples are equivalent to 1.

We also tested the activity of split-TF components containing TA sequences of varying levels of potency. In previous work, we demonstrated that substituting more potent transcriptional effectors into a drug-inducible TF system resulted in higher reporter activation levels at decreased drug concentrations^38^. Based on these previous results, we anticipated that variation of the TA sequence fused to HaloTag would enable a higher reporter response at lower levels of drug^38,39^. In the context of the present system, we predicted that elevated sensitivity in this manner may allow the induction of strong gene expression responses at low and intermediate drug concentrations, particularly at levels below the onset of “Hook effect”-mediated diminishment.

To test this possibility, we generated HaloTag fusions containing the potent TAs VP64-p65 and VP64-p65-RTA (VPR) (generating HaloTag-VP64-p65 and HaloTag-VPR) and compared these sequences alongside our original HaloTag-VP64. To evaluate their transcriptional activity, each HaloTag-TA fusion was co-expressed alongside Gal4_DB_-SNAP-tag in reporter cells (UAS H2B-mCherry). H2B-mCherry reporter expression levels from HaXS8 and drug-untreated cells were quantified via flow cytometry following overnight incubation (**Figure 2c**).

Consistent with our previous results, TAs of increased potency (HaloTag-VP64-p65 and HaloTag-VPR) resulted in stronger reporter expression levels at reduced drug concentrations (down to 1.6 nM) as compared to the original HaloTag-VP64. Notably, while HaloTag-VP64-p65 and HaloTag-VPR were able to elevate the drug sensitivity of transcriptional responses, expression of these more potent TAs did not appear to elevate levels of leaky/background transcription in the absence of drug. This suggests that un-dimerized HaloTag-TA proteins are unable to drive transcription without colocalization to DB domains, regardless of the strength of the TA. Encouraged by this finding, we also tested a variant of the system with multiple SNAP-tag domains, inspired by the SunTag system^40^ (**Supplemental Figure 4**). A Gal4_DB_-SNAP-tag-SNAP-tag protein, when paired with HaloTag-VP64, was able to drive a stronger transcriptional response than Gal4_DB_-SNAP-tag at the same drug concentrations, likely due to the possibility of recruiting two HaloTag-VP64 domains to a single Gal4_DB_-SNAP-tag-SNAP-tag domain.

While the two plasmid version of the SNAP-tag/HaloTag heterodimerizer system is useful in its ability to easily mix and match different DB-SNAP-tag constructs with different HaloTag-TA constructs, for ease of implementation and to ensure a similar ratio of both TF halves, we wanted to construct a version of the system where both halves were encoded on a single plasmid. For this system we utilized 2A “self-cleaving” peptides, which cause ribosome skipping during translation in eukaryotic cells, resulting in the “cleavage” of the polypeptide being translated^41^. The tandem P2A T2A was chosen due to its small size (40 aa), efficient “skipping” activity, and its generation of suitable levels of C-terminally positioned proteins within bicistronic fusions^41^.

Based on these considerations, we designed several constructs in the form of DB-SNAP-tag-P2A-T2A-HaloTag-TA (**Figure 2d**). Constructs were tested by expressing in mammalian cells with corresponding reporters, exposing to HaXS8, and analyzing by flow cytometry (**Figure 2e-g**). We found that for all DB and TA domain combinations, results were comparable or better to transfecting the system as two separate plasmids (**Figure 2f, g**). In particular, we noted that the background did not appear to be higher than the two plasmid system at both the protein and reporter level, suggesting that the P2A-T2A cleavage is very efficient (**Figure 2f, g, Supplemental Figures 5, 6**). Since output of both versions of the system are equivalent, a future user could choose which is more optimal for their specific experimental setup: the two plasmid system for modularity, or the single plasmid system for simplicity.

We do note that the covalent nature of HaXS8 dimerization limits the speed at which the split-TF system alone can be turned off. Reversibility of the system at a cellular level is possible, as when drug is removed the previously dimerized SNAP-tag and HaloTag fusion proteins will eventually be degraded, but the system more naturally lends itself to control of processes where reversibility is unnecessary or undesired. However, this limitation can be addressed by combining the HaXS8 dimerization system with other existing systems, such as the tetracycline inducible system^42^. By incorporating the TetR domain into a split TF (TetR-SNAP-tag, HaloTag-VP64), we were able to demonstrate that the addition of doxycycline was sufficient to force a faster “turn-off” of the system, both in the presence and after the removal of HaXS8 (**Supplemental Figure 7, Supplemental Video 1**).

In addition to controlling gene expression through a HaXS8-inducible split-TF system, we were also interested in controlling further protein behavior. Previously, the HaXS8-inducible system was shown to be able to activate the PI3K/mTOR pathway through localization of an inter-SH2 domain of p85 to the plasmid membrane^32^. We theorized that in addition to this, the HaXS8 dimerization system could be used to recover behavior of a split protein or activate proteins that require dimerization for activity. Furthermore, we specifically looked at biological processes where reversibility was unnecessary or undesired, as the SNAP-tag and HaloTag reactions with HaXS8 are covalent and thus irreversible. Examples of such processes include gene recombination, genome editing, and cell death.

The first system we considered for this was an inducible recombinase system. Having the ability to “knock-in” or “knock-out” genes in response to a small molecule drugs is a powerful tool (such as in mouse disease models^43^), so a variety of split recombinases controlled by CID systems have been constructed^16,44^. Inspired by these designs, we sought to determine whether our HaXS8-inducible system could be utilized in a similar manner.

Cre was chosen for a proof-of-concept experiment due to its widespread usage and well characterized reporter vectors^45^. A split site at aa270/271 was chosen due to its efficiency with the GIB, ABA, and rapamycin-inducible systems^16^, with the expectation that the split recombinase would only be active in the presence of drug (**Figure 3a**). Two variations of the system were constructed: a two plasmid system and a single plasmid system (**Figure 3b**). To determine whether the HaXS8-inducible split Cre system was functional, these constructs were transfected into mammalian cells stably expressing a dsRed to GFP Cre stoplight reporter^45^ and exposed to varying levels of drug. Analysis via flow cytometry demonstrated that both versions of the system responded in a concentration dependent manner to HaXS8, with the P2A T2A single plasmid system being most effective (**Figure 3c, d**). Maximal reporter activity was observed at a concentration of 200nM for both the two plasmid system (at 2 different ratios) and the single plasmid system. It is relevant to note that background of the single plasmid system was comparable to that of a miRFP670 only control (~3% GFP positive). This suggests that our split Cre(aa1-270)-SNAP-tag and HaloTag-Cre(aa271-343) are not spontaneously recovering activity without the HaXS8 dimerizer present, and that cleavage of our P2A T2A sequence is extremely efficient. Any failure of cleavage of the P2A T2A sequence would result in a fusion containing both split Cre halves, so lack of background is a promising sign that failure of cleavage is extremely rare.

**Figure 3.**
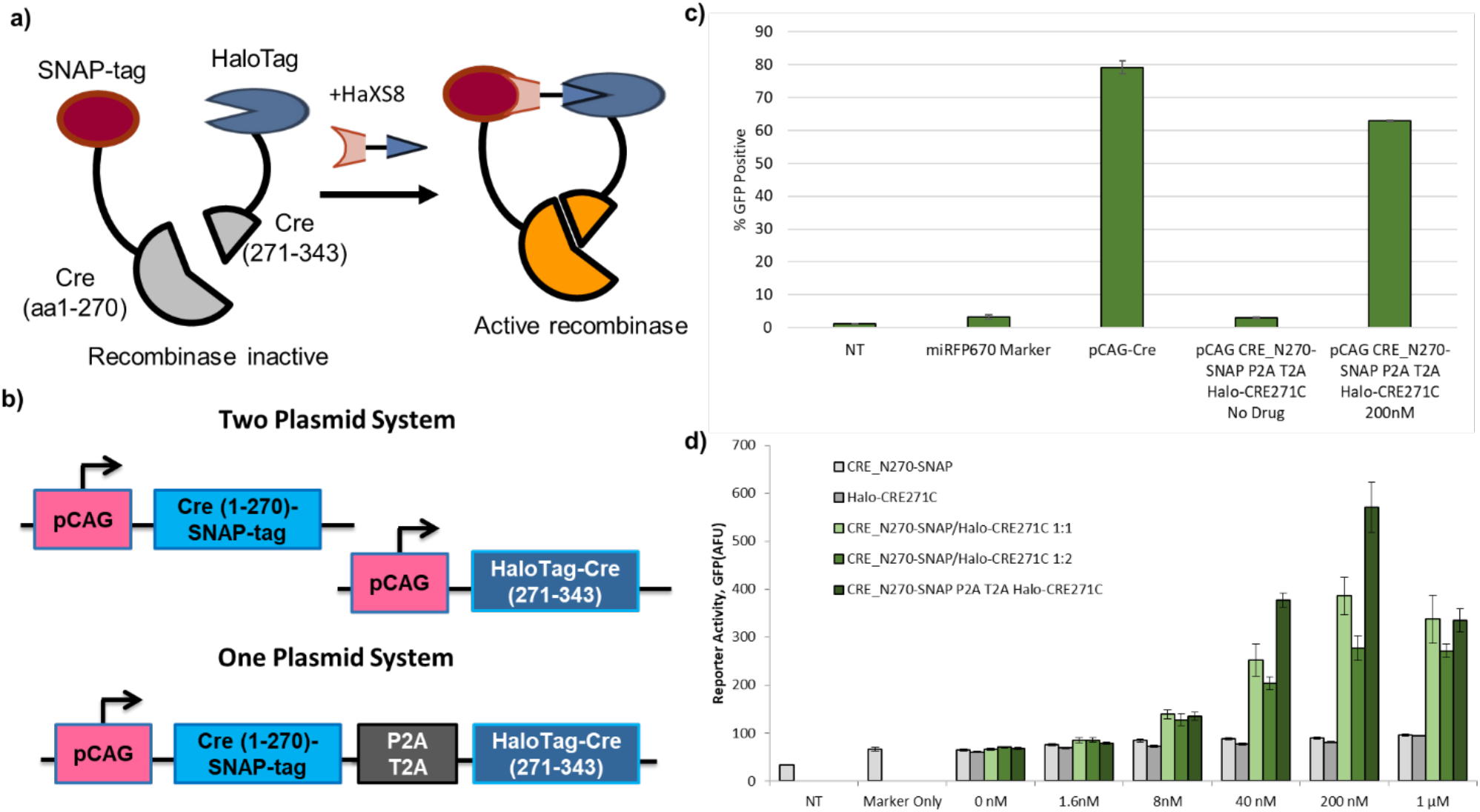
Design of a HaXS8-inducible recombinase system. **a)** Protein schematic for the two parts of the HaXS8-inducible recombinase system. Cre(1-270) is fused to a SNAP-tag domain, and Cre(271-343) is fused to a HaloTag domain. Addition of HaXS8 to the system allows for heterodimerization and thus active recombinase. **b)** Schematic showing the plasmid design for two variations of a HaXS8-inducible split-Cre recombinase system. **c)** Percent GFP positive cells as determined by flow cytometry in Hek293FT reporter cells (dsRed->GFP Cre stoplight) transiently transfected with the indicated plasmids and treated with varying concentrations of HaXS8. Reporter cells yield GFP expression upon site-specific recombination. Cells expressing GFP equal to or higher than the top 1% of a non-transfected control are considered “GFP positive.” Marker refers to a miRFP670 transfection marker. Geometric means are displayed as mean ± s.d., as determined by three transfected cell cultures. **d)** GFP fluorescence as determined by flow cytometry in Hek293FT reporter cells (dsRed->GFP Cre stoplight) transiently transfected with the indicated plasmids and treated with varying concentrations of HaXS8. CRE_N270-SNAP refers to only the Cre(aa1-270)-SNAP-tag encoding part, and Halo-CRE271C refers to HaloTag-Cre(aa271-343). CRE_N270-SNAP and Halo-CRE271C were transfected at both a 1:1 and 1:2 ratio. CRE_N270-SNAP P2A T2A Halo-CRE271C refers to Cre(aa1-270)-SNAP-tag P2A T2A HaloTag-Cre(aa271-343). Reporter cells yield GFP expression upon site-specific recombination. Marker refers to a miRFP670 transfection marker. Geometric means are displayed as mean ± s.d., as determined by three transfected cell cultures.

In addition to controlling behavior of split-proteins, CID systems have also been used to drive a variety of cell behaviors through protein interaction, including ones relevant to cellular therapeutics such as apoptosis. The inducible caspase-9 (iCasp9) system has been utilized in studies as a “safety switch” for T-cell therapies^46^, and more recently, a rapamycin-based caspase-9 system has also been developed^7^. Based on these systems, we hypothesized that we could create a HaXS8-inducible caspase-9 dimerization system. In the absence of HaXS8, SNAP-tag-caspase-9 and HaloTag-caspase-9 fusion proteins would remain separate. However, in the presence of HaXS8, the two proteins would dimerizer, leading to apoptosis (**Figure 4a**). We designed a vector encoding HaloTag-caspase-9 and SNAP-tag-caspase-9, each driven from opposing arms of a bidirectional CMV promoter (**Figure 4b**). To test the system, we observed mammalian cells expressing our construct and exposed them to HaXS8. Fluorescent microscopy showed that tdTomato fluorescence was dependent on HaXS8 concentration, with fluorescence being the lowest at 1μM and increasing at 5μM, matching dimerization behavior previously observed in the HaXS8-inducible TF system (**Figure 4c**). Such drug dependence was not observed in a SNAP-tag-caspase-9 only control, which expressed the same vector lacking the HaloTag-caspase-9 sequence (**Figure 4d)**. This same trend was confirmed through flow cytometry (**Figure 4e, f**). However, we note that a subset of transfected cells do remain alive, even when exposed to drug. From analysis of tdTomato expression via imaging and flow cytometry, we hypothesize that this is due to the highly expressing subset of the population being more sensitive to drug. Increased delivery via viral transduction or further sorting to select a highly expressing population is likely necessary for a population that is fully killed off by the addition of HaXS8. Such necessity of high expression in caspase-9 inducible systems has previously been observed by other studies^7,47^.

**Figure 4.**
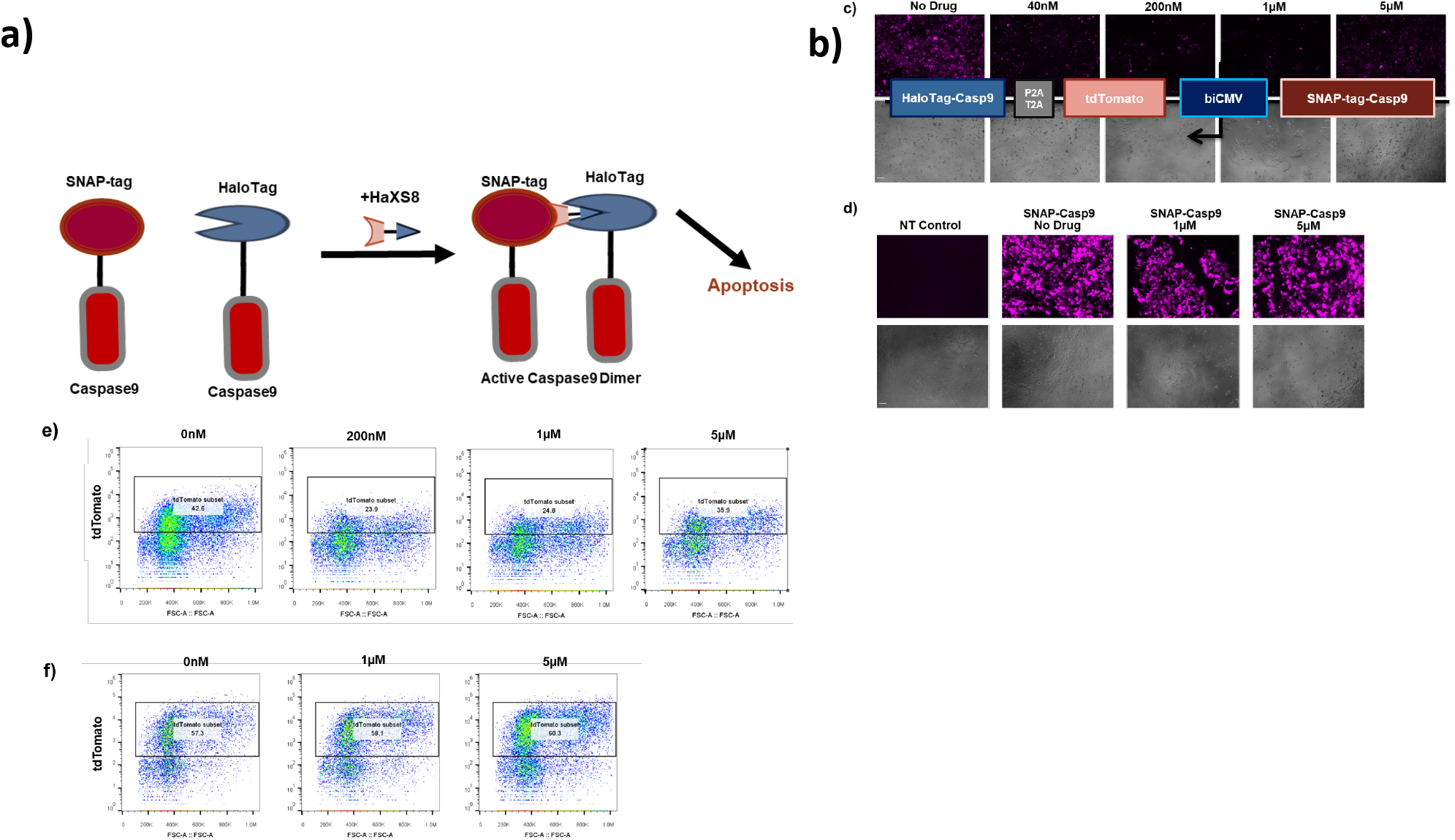
Design of a HaXS8-inducible apoptosis system. **a)** Schematic demonstrating the behavior for the HaXS8-inducible apoptosis system. In the absence of HaXS8, SNAP-tag-caspase-9 (SNAP-Casp9) and HaloTag-caspase-9 (Halo-Casp9) remain separate. In the presence of HaXS8, the two domains dimerizer, activating downstream effects which result in apoptosis. **b)** Schematic showing the plasmid design for the HaXS8-inducible Casp9 dimerization system. A bidirectional CMV promoter (biCMV) drives SNAP-tag-caspase-9 on one side, and tdTomato-P2A-T2A-HaloTag-caspase-9 on the other. **c)** Fluorescence images of HEK 293FT cells transiently transfected with the vector shown in **(b)**, treated with varying concentrations of HaXS8. Upper row shows tdTomato fluorescence. Lower row shows brightfield. Scale bar is 100 μm. **d)** Fluorescence images of non-transfected HEK 293FT cells or HEK293FT cells transiently transfected with a vector encoding only tdTomato and SNAP-tag-caspase-9, treated with varying concentrations of HaXS8. Upper row shows tdTomato fluorescence. Lower row shows brightfield. Scale bar is 100 μm. **e)** Representative plots for the HaXS8-inducible Casp9 dimerization system with tdTomato shown against the FSC-A. Boxed population shows a cell population highly expressing tdTomato, and thus highly expressing the HaXS8-inducible system. Cells are gated to exclude non-cell debris. A representative experiment is shown. **f)** Representative plots for a SNAP-tag-Casp9 control with tdTomato shown against the FSC-A. Boxed population shows a cell population highly expressing tdTomato, and thus highly expressing the transfected vector. Cells are gated to exclude noncell debris. A representative experiment is shown. Cells were transfected ~20 hours before HaXS8 addition, which was ~30 hours before imaging.

## DISCUSSION

In this study, we described additional applications of a small-molecule heterodimerizer system based on the covalent dimerization of the SNAP-tag and HaloTag domains with HaXS8. We demonstrated that in addition to its previous usage for protein localization^32^, the HaXS8 system can also be used as a general system to control gene expression via split TFs, protein activity via split Cre, and cellular behavior via Casp9 dimerization.

The split-TF system was shown to be drug-dependent to various concentrations of HaXS8, both in dimerizing the two halves of the split-TF system and in activating expression of a reporter gene. Dimerization was shown to be sensitive and fast, with immunoblotting showing intact dimerized protein at concentrations as low as 1.6nM and in as little time as 24 minutes. Various DB and TA domains were tested, demonstrating the modularity of the system, and the number of SNAP-tag domains were modulated to control stoichiometry of HaloTag-TA units that could bind individual DB-(SNAP-tag)_n_ domains. The system was also converted to a single plasmid system for easier implementation. We anticipate that other activating or repressing domains could be combined with the system to allow additional sophistication in HaXS8-mediated gene expression control.

HaXS8 was also used to control split Cre recombinase, demonstrating drug-dependent response of a GFP reporter. This HaXS8-inducible Cre recombinase was tested in both a 2-vector and single vector system, and the single vector system seemed marginally more efficient. Notably, background remained low for this single vector design. This initial design could be further extended for use with other Cre split-sites or other split recombinases for further utility.

A HaXS8-inducible Casp9 dimerization system was also developed by creating SNAP-tag-Casp9 and HaloTag-Casp9 fusion proteins. Albeit most effective in a population of cells highly expressing the construct, we were able to observe what appears to be a drug-dependent decrease of cells in this population, suggesting that apoptosis was occurring in response to drug. This drug dependency was not observed in cells only expressing the SNAP-tag-Casp9 half of the system, further supporting our claim that the HaXS8 dimerization of SNAP-tag-Casp9 and HaloTag-Casp9 is inducing apoptosis. Selecting for higher expression through cell sorting or generating a clonal line is likely to help us further confirm the system, as other inducible-caspase-9 systems also require high levels of expression to function^7,47^.

A variety of potential future directions exist for the system. One of the most straightforward would be to test the system with the photocleavable MeNV-HaXS, which is cleaved in response to UV light^48^. This would allow for reversibility of the system, which would potentially be useful in turning off activation from our HaXS8-inducible TF technique. Furthermore, since the cleavage is in response to light, this deactivation would have spatiotemporal control. Combining the HaXS8-inducible system with other drug-inducible systems opens up additional directions for controlling more complex behavior or creating multi-input genetic circuits. Since the system has been demonstrated to be orthogonal to both the rapamycin inducible system^32^ and the LInC system^38^ (**Supplemental Figure 8, 9**), there are at least 2 potential orthogonal small molecules that could be applied. Due to the bioorthagonality of SNAP-tag, HaloTag, and their ligands^49^, it seems likely that other CID systems, such as the GIB and ABA inducible systems, would be orthogonal and applicable as well.

The applications presented here demonstrate that the HaXS8-inducible system can be used to install drug sensitivity into a variety of proteins of synthetic biology utility. As SNAP-tag, HaloTag, and their substrates are biorthogonal, they are not expected to interfere with cellular behavior. Given the conventional wisdom that *sola dosis facit venenum* (“the dose makes the poison”), minimization of required drug doses is a desirable feature of any inducible system. In our split-TF work, we showed that HaXS8 doses could be lowered without sacrificing transcriptional response levels by using more potent TA domains. Use of SNAP- and Halo tags also affords certain advantages in their laboratory analysis and implementation, due to commercially available antibodies and their various reactive ligands based on fluorescent dyes, biotin, and functionalized beads. The fact that the heterodimerization of SNAP-tag and HaloTag with HaXS8 is covalent also makes dimerization in a target system directly quantifiable, as opposed to having to depend on output from a further downstream process. While to our knowledge there has been no use of HaXS8 in *in vivo* studies, HaloTag^50^ and SNAP-tag^51^ have both been demonstrated to work *in vivo*. The results shown here establish a variety of uses with HaXS8 *in vitro*, and it stands to reason that the system could be extended to *in vivo* studies. We hope that the new applications of the HaXS8-inducible dimerization system described in this work will add to the existing protein dimerization toolkit for gene expression and protein control, in addition to HaXS8’s already existing context as a protein localization tool. To facilitate such work, plasmid DNAs and full sequence information for the described constructs have been deposited to AddGene, with note that HaXS8 can currently be purchased via vendor suppliers (as of the date of this publication; see Materials and Methods).

## METHODS

### DNA constructs

Standard cloning procedures were used in the generation of all DNA constructs. Plasmid DNA and detailed sequence information for described constructs were deposited to AddGene upon acceptance of this publication.

### Small molecule inducers

HaXS8 was from Tocris (Cat. No. 4991). MK-5172 (grazoprevir) was from MedChemExpress. Small molecule-inducer stocks were dissolved in DMSO at a concentration of 10 mM and diluted into cell culture media at the indicated working concentrations.

### Mammalian cell culture

All mammalian cell lines were cultured in a humidified incubator maintained at 37 °C with 5% CO2. HEK293FT cells (Thermo Fisher) were cultured in DMEM with 10% FBS and supplemented with nonessential amino acids (Life Technologies) and Glutamax (Life Technologies).

### DNA transfections

DNA transfections were carried out with Lipofectamine 3000 Reagent (Thermo Fisher) according to the manufacturer’s instructions. For imaging experiments, cells were seeded in dishes or well plates containing coverslip bottoms either coated with bovine plasma fibronectin (Product #F1141, Sigma-Aldrich) or treated for cell adherence by the manufacturer (poly-D-lysine by MatTek, or ibiTreat by ibidi).

### Antibodies

The following primary antibodies were used: mouse anti-HA-HRP (Cell Signaling; 6E2, #2999; 1:1,000 dilution for western blotting), rabbit anti-SNAP-tag (NEB; P9310; 1:1,000 dilution for western blotting), rabbit anti-T7-tag (Cell Signaling; D9E1X, #13246; 1:1000 dilution for western blotting), mouse anti-TetR (Clontech; 9G9; 1:1000 dilution for western blotting), mouse anti-GAPDH (Thermo Fisher; GA1R; 1:3,000 dilution for western blotting), rabbit anti-GAPDH (Sigma-Aldrich; G9545; 1:3,000 dilution for western blotting), and rat anti-GAPDH (Biolegend; W17079A; 1:10,000 dilution for western blotting). The following secondary antibodies were used: anti-mouse HRP conjugate (Cell Signaling; 7076; 1:3,000 dilution), and anti-rabbit HRP conjugate (Bio-Rad; 170-6515; 1:3,000 dilution).

### Preparation of cell lysates for immunoblotting

Cell lysates used in immunoblotting analyses were prepared by direct lysis of drug-treated and untreated cells in 1× LDS–PAGE loading buffer (Thermo Fisher) after removal of cell culture media. Viscous solutions were formed after addition of the lysis reagent, and were clarified through sonication followed by centrifugation. The lysates were subsequently analyzed by standard immunoblotting procedures and probed using the antibodies listed above at the indicated dilutions. Detection of the labeled antigens was carried by chemiluminescence via the SuperSignal West Pico PLUS Chemiluminescent Substrate (Pierce).

### Image acquisition and analysis

Cells were imaged by epifluorescence microscopy in imaging-compatible vessels containing glass coverslip bottoms (MatTek) or optically clear plastic bottoms (ibidi). During imaging, cells were maintained in PBS, standard culture media, or FluoroBrite DMEM (Thermo Fisher). Images were acquired with ZEN imaging software (Zeiss). Image files were processed with the ImageJ-based image analysis package Fiji. The images were contrasted uniformly across experiments.

### Flow cytometry

Cells analyzed by flow cytometry were gated for living cells by scatter detection. The geometric mean measured reporter fluorescence levels were reported in arbitrary fluorescence units (AFU). Reporter activation analyses were carried out with stable single clones or cells transiently transfected with DNA encoding the analyzed TF or enzyme (as indicated in the figure captions). For analyses carried out with transiently expressing cells, plasmid DNA encoding a constitutively expressed fluorescent protein marker was co-delivered at the time of transfection and used to identify positively transfected cell populations. Briefly, transfected cells were gated to the top 1% of marker fluorescence of nontransfected control cells under the same condition. Transient expression experiments carried out with a portion of the turn-on TFs were gated via detection of an mCherry marker that was expressed via an IRES sequence on the TF-encoding plasmid. mCherry marker was expressed on either one of the two halves of the two-part system, or on in the single vector variation. Transfected cells were incubated for 24–48 h after transfection in either the presence or the absence of the indicated small molecules before being analyzed with an Attune NxT flow cytometer (Thermo Fisher). For the analyses in which DB-SNAP-tag and HaloTag-TA constructs are used together, DNA mixtures containing a 1:1 ratio were used unless otherwise specified.

