## Supplementary Information v1 for "SNAP-tag and HaloTag fused proteins for HaSX8-inducible control over synthetic biological functions in engineered mammalian cells"

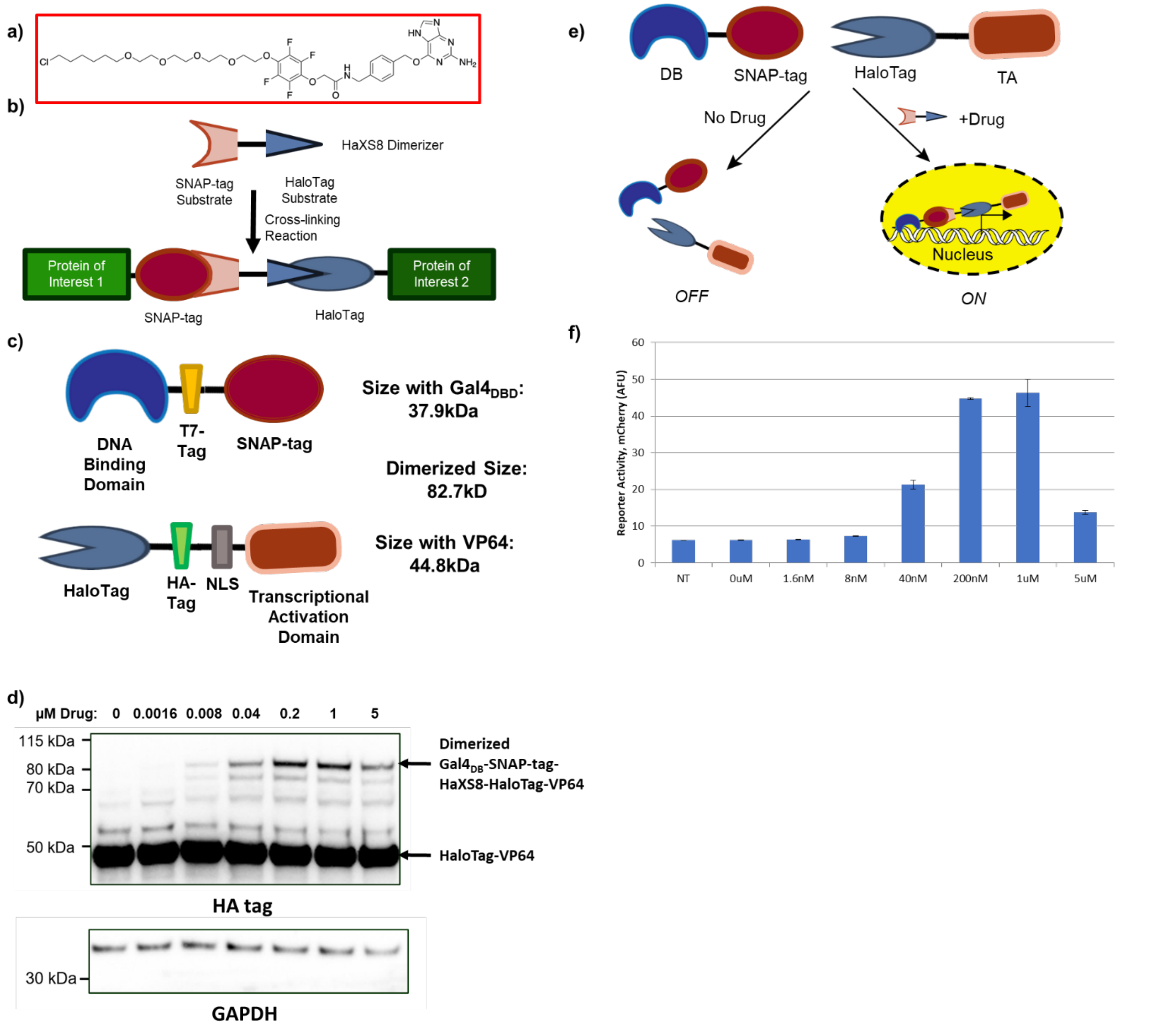


**Figure 1.** ***Design of a HaXS8-inducible TF system.*** **a)** Chemical structure of the HaXS8 molecule. **b)** HaXS8 consists of a SNAP-tag substrate and a HaloTag substrate connected by a linking module. HaXS8 is able to heterodimerize proteins of interest fused to SNAP-tags and HaloTags. **c)** Schematic showing the structure of the HaXS8-inducible TF system, as well as tag locations for immunoblotting. **d)** Western blot showing accumulation of full-length, dimerized Gal4$_{DB}$-SNAP-tag-HaXS8-HaloTag-VP64 (anti-HA; 82.7 kDa) in response to HaXS8. Undimerized HaloTag-VP64 (anti-HA; 44.8 kDa) is also observed. GAPDH (anti-GAPDH; 37 kDa) was used as a loading control. **e)** When no drug is present, DB-SNAP (DNA-Binding Domain-SNAP-tag) and Halo-TA (HaloTag-Transcriptional Activation Domain) remain separate, and do not act as a functional TF. When HaXS8 is added, the two halves are able to heterodimerize, resulting in a functional TF. This functional TF allows transcription of a target gene that corresponds with concentration of drug. **f)** H2B-mCherry fluorescence as determined by flow cytometry in Hek293FT reporter cells (UAS H2B-mCherry) transiently transfected with plasmids encoding Gal4$_{DB}$-SNAP-tag and HaloTag-VP64 at a 1:1 ratio and treated with varying concentrations of HaXS8. Geometric means are displayed as mean ± s.d., as determined by three transfected cell cultures.

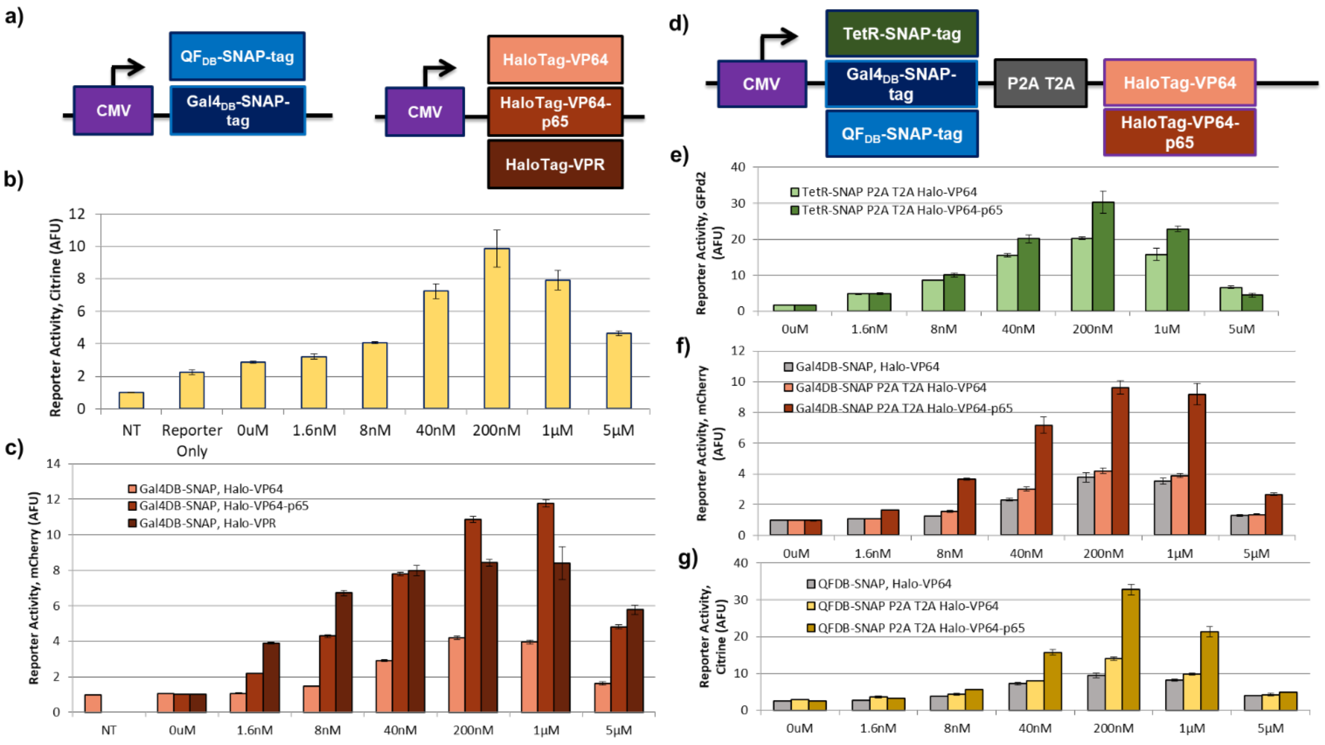


**Figure 2.** ***Designs with varying DB and TA domains for a HaXS8-inducible "Turn-on" system.*** **a)** Schematic showing the plasmid schematics for the two halves of the HaXS8-inducible TF: DB-SNAP (DNA-Binding Domain-SNAP-tag) and (Halo-TA) HaloTag-Transcriptional Activation Domain. **b)** H2B-citrine fluorescence as determined by flow cytometry in Hek293FT cells transiently transfected with a QUAS H2B-citrine reporter and plasmids encoding $QF_{DB}$-SNAP-tag and HaloTag-VP64 at a 1:1 ratio, treated with varying concentrations of HaXS8. **c)** H2B-mCherry fluorescence as determined by flow cytometry in Hek293FT reporter cells (UAS H2B-mCherry) transiently transfected with plasmids encoding $Gal4_{DB}$-SNAP-tag and varying HaloTag-TAs at a 1:1 ratio and treated with varying concentrations of HaXS8. **d)** ***Single vector design for a HaXS8-inducible "Turn-on" system.*** Schematic showing the design of a single vector encoding both halves of the HaXS8-inducible "turn-on" system. Both halves of the HaXS8-inducible TF, DB-SNAP (DNA-Binding Domain-SNAP-tag) and Halo-TA (HaloTag-Transcriptional Activation Domain), are encoded on one plasmid, separated by a P2A T2A sequence. **e)** GFPd2 fluorescence as determined by flow cytometry in Hek293FT reporter cells (TRE3G GFPd2) transiently transfected with a plasmid encoding either TetR-SNAP-tag P2A T2A HaloTag-VP64 or TetR-SNAP-tag P2A T2A HaloTag-VP64-p65, treated with varying concentrations of HaXS8. **f)** H2B-mCherry fluorescence as determined by flow cytometry in Hek293FT reporter cells (UAS H2B-mCherry) transiently transfected with a plasmid encoding either $Gal4_{DB}$-SNAP-tag P2A T2A HaloTag-VP64 or $Gal4_{DB}$-SNAP-tag P2A T2A HaloTag-VP64-p65, treated with varying concentrations of HaXS8. For comparison, Hek293FT reporter cells (UAS H2B-mCherry) transiently transfected with plasmids encoding $Gal4_{DB}$-SNAP-tag and HaloTag-VP64 at a 1:1 ratio, treated with varying concentrations of HaXS8, are also shown. **g)** H2B-citrine fluorescence as determined by flow cytometry in Hek293FT cells transiently transfected with a QUAS H2B-citrine reporter and a plasmid encoding either $QF_{DB}$-SNAP-tag P2A T2A HaloTag-VP64 or $QF_{DB}$-SNAP-tag P2A T2A HaloTag-VP64-p65, treated with varying concentrations of HaXS8. For comparison, Hek293FT cells transiently transfected with a QUAS H2B-citrine reporter and plasmids encoding $QF_{DB}$-SNAP-tag and HaloTag-VP64 at a 1:1 ratio, treated with varying concentrations of HaXS8, are also shown. Geometric means are displayed as mean ± s.d., as determined by three transfected cell cultures. Data is normalized such that NT samples are equivalent to 1.

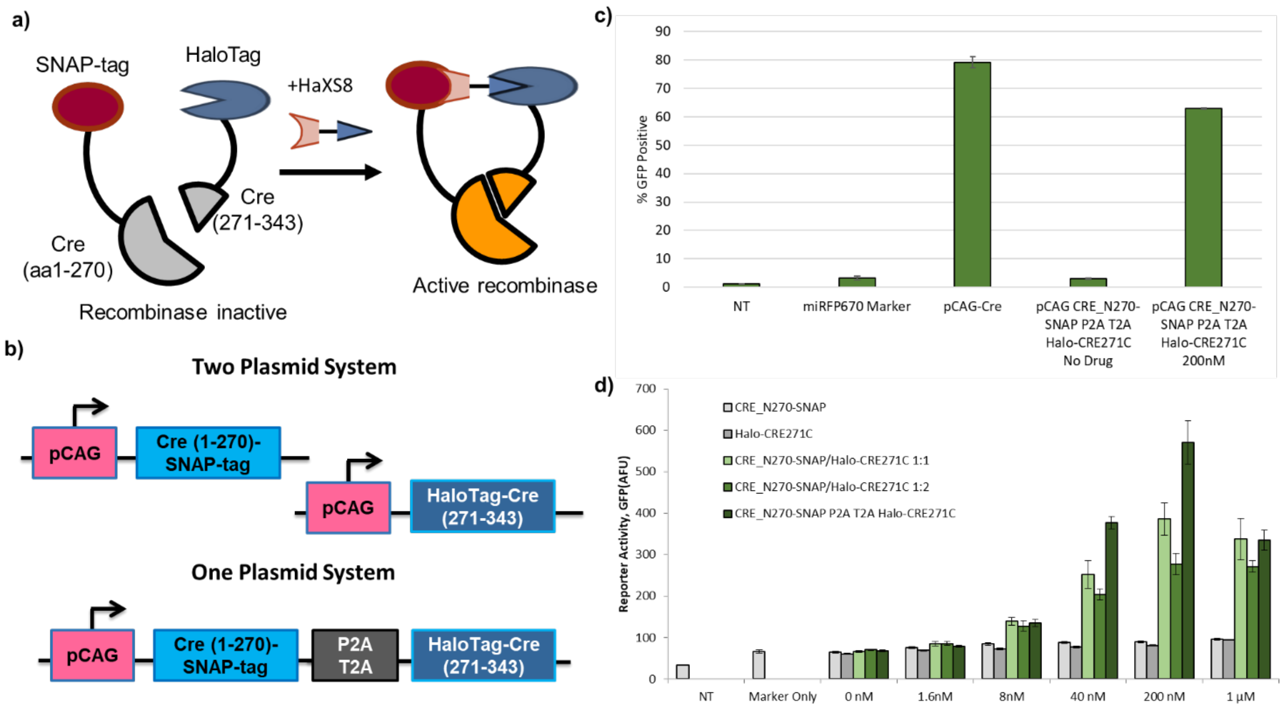


**Figure 3.** ***Design of a HaXS8-inducible recombinase system.*** **a)** Protein schematic for the two parts of the HaXS8-inducible recombinase system. Cre(1-270) is fused to a SNAP-tag domain, and Cre(271-343) is fused to a HaloTag domain. Addition of HaXS8 to the system allows for heterodimerization and thus active recombinase. **b)** Schematic showing the plasmid design for two variations of a HaXS8-inducible split-Cre recombinase system. **c)** Percent GFP positive cells as determined by flow cytometry in Hek293FT reporter cells (dsRed->GFP Cre stoplight) transiently transfected with the indicated plasmids and treated with varying concentrations of HaXS8. Reporter cells yield GFP expression upon site-specific recombination. Cells expressing GFP equal to or higher than the top 1% of a non-transfected control are considered "GFP positive." Marker refers to a miRFP670 transfection marker. Geometric means are displayed as mean ± s.d., as determined by three transfected cell cultures. **d)** GFP fluorescence as determined by flow cytometry in Hek293FT reporter cells (dsRed->GFP Cre stoplight) transiently transfected with the indicated plasmids and treated with varying concentrations of HaXS8. CRE_N270-SNAP refers to only the Cre(aa1-270)-SNAP-tag encoding part, and Halo-CRE271C refers to HaloTag-Cre(aa271-343). CRE_N270-SNAP and Halo-CRE271C were transfected at both a 1:1 and 1:2 ratio. CRE_N270-SNAP P2A T2A Halo-CRE271C refers to Cre(aa1-270)-SNAP-tag P2A T2A HaloTag-Cre(aa271-343). Reporter cells yield GFP expression upon site-specific recombination. Marker refers to a miRFP670 transfection marker. Geometric means are displayed as mean ± s.d., as determined by three transfected cell cultures.

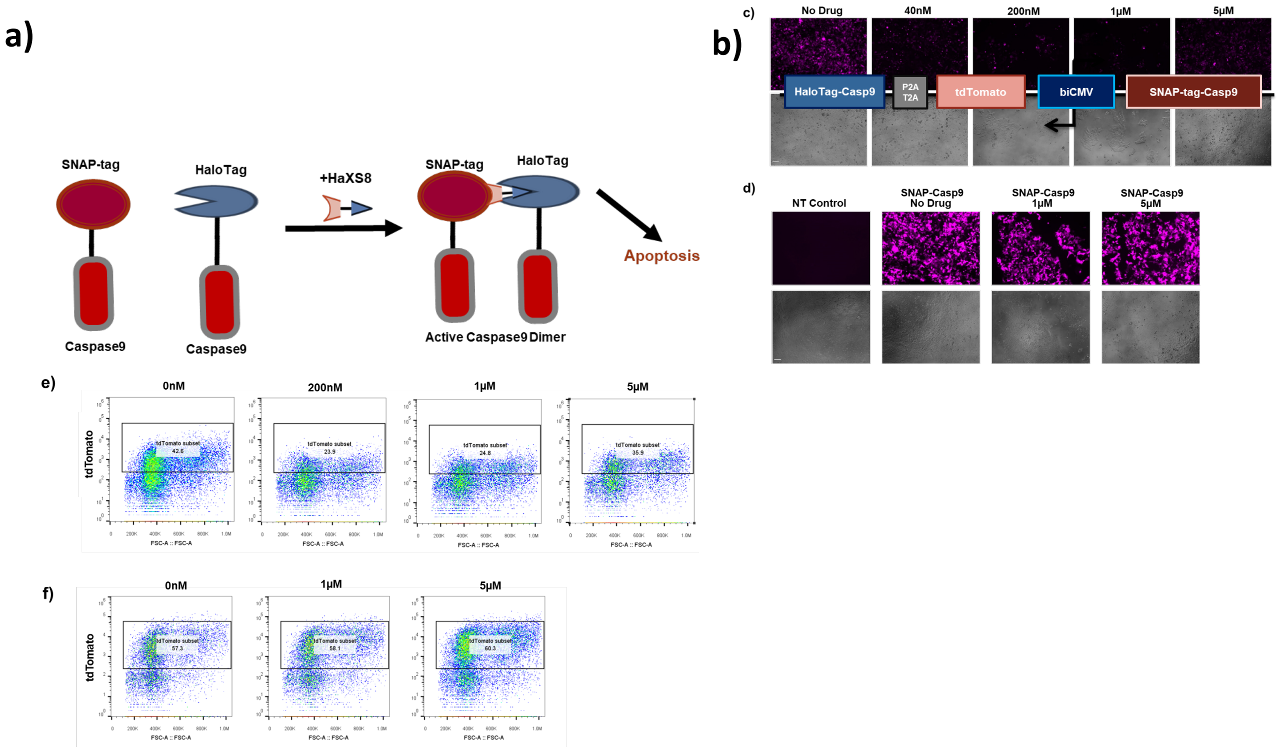


**Figure 4.** ***Design of a HaXS8-inducible apoptosis system.*** **a)** Schematic demonstrating the behavior for the HaXS8-inducible apoptosis system. In the absence of HaXS8, SNAP-tag-caspase-9 (SNAP-Casp9) and HaloTag-caspase-9 (Halo-Casp9) remain separate. In the presence of HaXS8, the two domains dimerizer, activating downstream effects which result in apoptosis. **b)** Schematic showing the plasmid design for the HaXS8-inducible Casp9 dimerization system. A bidirectional CMV promoter (biCMV) drives SNAP-tag-caspase-9 on one side, and tdTomato-P2A-T2A-HaloTag-caspase-9 on the other. **c)** Fluorescence images of HEK 293FT cells transiently transfected with the vector shown in **(b)**, treated with varying concentrations of HaXS8. Upper row shows tdTomato fluorescence. Lower row shows brightfield. Scale bar is 100 μm. **d)** Fluorescence images of non-transfected HEK 293FT cells or HEK293FT cells transiently transfected with a vector encoding only tdTomato and SNAP-tag-caspase-9, treated with varying concentrations of HaXS8. Upper row shows tdTomato fluorescence. Lower row shows brightfield. Scale bar is 100 μm. **e)** Representative plots for the HaXS8-inducible Casp9 dimerization system with tdTomato shown against the FSC-A. Boxed population shows a cell population highly expressing tdTomato, and thus highly expressing the HaXS8-inducible system. Cells are gated to exclude non-cell debris. A representative experiment is shown. **f)** Representative plots for a SNAP-tag-Casp9 control with tdTomato shown against the FSC-A. Boxed population shows a cell population highly expressing tdTomato, and thus highly expressing the transfected vector. Cells are gated to exclude non-cell debris. A representative experiment is shown. Cells were transfected ~20 hours before HaXS8 addition, which was ~30 hours before imaging.
